# Antiviral activity of anisomycin against chikungunya virus

**DOI:** 10.64898/2026.06.19.733322

**Authors:** Sho Kawashima, Akino Emi, Fumiyo Ogawa, Shoichi Sakaguchi, Taku Ogawa, Hong Wu, Hirotaka Ebina, Youichi Suzuki, Takashi Nakano

**Author notes:** These authors contributed equally to this work (Sho Kawashima, Akino Emi). Corresponding author (YS).

## Abstract

Chikungunya virus (CHIKV) is a globally prevalent arbovirus transmitted by *Aedes* mosquitoes, which causes acute fever accompanied by debilitating joint pain that can persist for extended periods. Despite the significant public health impact and an increasing incidence worldwide, antiviral treatment targeting CHIKV has not been clinically approved. In this study, we screened compounds using a newly developed In-Cell ELISA-based assay and CHIKV Indian Ocean Lineage (IOL) and found that an antibiotic derived from *Streptomyces* bacteria, anisomycin, potentially inhibited CHIKV. The selectivity index of anisomycin was favorable for anti-CHIKV activity, with 50% effective concentration (EC_50_) of 200 pM and 50% cytotoxic concentration (CC_50_) of 390 nM in Vero cells. This robust inhibitory activity against CHIKV was confirmed in a human cell line and against a CHIKV East/Central/South African (ECSA) lineage. These effects of anisomycin were apparently independent of its functions as a translation inhibitor and mitogen-activated protein kinase (MAPK) pathway stimulator. These findings, together with the finding that anisomycin suppressed the production of infectious CHIKV virions, suggested that anisomycin inhibits CHIKV *via* a distinct mechanism. Further mechanistic insights were gained through genetic analyses of anisomycin-resistant mutants, which revealed that a single amino acid substitution (G117R) in the macrodomain of CHIKV nsP3 confers resistance to anisomycin. Importantly, anisomycin reduced footpad swelling and viremia in mice during the early days of CHIKV infection, indicating its therapeutic potential. Given its inhibitory activity against other arboviruses, our study positions anisomycin as a promising lead inhibitor for the future development of broad-spectrum antiviral drugs, including CHIKV.

**Author summary:** Chikungunya fever (CHIKF) is a mosquito-borne disease caused by the chikungunya virus (CHIKV) and is characterized by fever, rash, and arthralgia. Although most persons infected with CHIKV recover within days, joint pain and severe complications can persist. However, the management of CHIKF is limited to symptom relief, and specific antiviral treatments are not available. Our study focused on identifying potential inhibitors of CHIKV infection. We found that the natural alkaloid, anisomycin, inhibited CHIKV replication in cultured cells *in vitro* using a novel screening assay and a chemical compound library. Interestingly, the mechanism by which anisomycin blocks CHIKV infection likely differs from its currently known effects, suggesting a distinct mode of inhibition. We also identified an amino acid change in a nonstructural protein that conferred resistance to anisomycin, providing insights into a viral target of anisomycin. Importantly, anisomycin reduced disease symptoms and viremia in mouse models of CHIKV *in vivo*. Because anisomycin inhibits other mosquito-borne viruses, our findings suggest that it could serve as a basis for developing broad-acting antiviral drugs.

## Introduction

Chikungunya virus (CHIKV) in the genus *Alphavirus* of the *Togaviridae* family is categorized as an enveloped arbovirus because it is transmitted to humans bitten by infected *Aedes aegypti* and *Aedes albopictus* mosquitoes [1]. Symptomatic chikungunya fever (CHIKF) caused by CHIKV is a febrile illness characterized by headache, fatigue, myalgia, nausea, skin rash, and symmetrical polyarthralgia involving distal joints such as the wrists, ankles, and toes. While CHIKV infection generally presents with a symptomatic illness resembling dengue fever, the symptoms mostly resolve within 10 days, and the mortality rate is low. However, the risk of severe complications, including encephalitis, cardiovascular disorders, renal failure, hepatitis, and myocarditis, can increase, particularly in vulnerable populations such as children, elderly persons, and individuals with preexisting conditions such as hypertension, diabetes, and heart disease [2,3]. Chronic arthralgia or arthritis persists in a substantial proportion of patients with CHIKV, and this is suggested to be linked to immune-mediated mechanisms rather than persistent CHIKV infection [4,5]. The U.S. Food and Drug Administration (FDA) recently approved the live attenuated vaccine, VLA1553 (IXCHIQ), which is effective as a prophylactic agent against the onset of CHIKF [6]. In contrast, treatment of patients infected with CHIKV is currently limited to supportive anti-inflammatory drugs and analgesics because an antiviral drug that specifically targets CHIKV is not clinically available [7].

The CHIKV particle contains a single-stranded positive-sense RNA genome of approximately 12 kb that encodes four nonstructural proteins (nsP1, nsP2, nsP3, and nsP4) on the 5’ side and five structural proteins (C, E3, E2, 6k, and E1) on the 3’ side. After binding to cell-surface receptors *via* envelope glycoproteins, CHIKV enters target cells by endocytosis, Thereafter, released viral RNA serves as a template to synthesize nonstructural proteins that form a replicase complex that produces full-length genomic and subgenomic (26S) RNAs. Structural proteins are translated from 26S RNA, proteolytically processed, and then nucleocapsid and envelope proteins assemble, and virions are released by budding through the plasma membrane [1,2].

The CHIKV lineages classified as West African (WA), East/Central/South African (ECSA), and Asian are based primarily on genetic variations in the E1 glycoprotein [8]. Although CHIKV has historically been associated with endemic transmission and sporadic outbreaks in Africa and Asia, it attracted global attention after a large-scale outbreak in the Indian Ocean during 2004. This was driven in part by the emergence of the Indian Ocean lineage (IOL) from the ECSA lineage. An important feature of IOL is its Ala-to-Val substitution at position 226 of E1, which enhances viral fitness in *Aedes albopictus* and facilitates transmission in tropical and temperate regions [8–10]. Furthermore, the Asian lineage continues to spread and has been implicated in outbreaks in India and South America. Accordingly, CHIKV has become a significant global public health threat as sustained transmission has persisted in more than 100 countries [11]. Therefore, an effective treatment for CHIKV infection is urgently required.

In an effort to develop anti-CHIKV drugs, we aimed to identify promising compounds that could inhibit CHIKV infection. Screening a commercial library of chemicals revealed that the natural alkaloid anisomycin potently inhibits CHIKV. The selectivity index of anisomycin in cultured cells was favorable, and the production of infectious virions was hampered. Anisomycin alleviated swollen feet induced by CHIKV and reduced the viral burden in model mice *in vivo*, indicating that it protects against CHIKV infection.

## Methods

### Cells and viruses

All cell cultures were maintained in appropriate culture medium supplemented with 10% fetal bovine serum (FBS) and antibiotics (100 units/ml penicillin and 100 μg/ml streptomycin) at 37°C under a 5% CO_2_. Vero (green monkey kidney cells, provided by the National Institute of Infectious Diseases, Japan) were cultured in Eagle’s minimal essential medium (MEM, Nacalai Tesque). Huh7 (human hepatoma cell, Japanese Collection of Research Bioresources) was cultured using Dulbecco’s modified Eagle’s medium (DMEM) with low (1 g/L) glucose (Nacalai Tesque). 293T cells (human embryonic kidney cells; Clontech) were cultured in DMEM with high (4.5 g/L) glucose (Nacalai Tesque).

Infectious CHIKV clone was produced in Vero cells by transfection with a plasmid DNA encoding the full-length viral genome (pCMV-SL11131 and pCMV-Ross) [12]. Infectious titers of CHIKV were measured by plaque assays using Vero cells as described previously [12].

### In-Cell ELISA

Vero cells (2 × 10^4^ cells/well) were seeded in white 96-well flat-bottom plates (SPL Life Sciences Co., Ltd.), then infected one day later with CHIKV SL11131 at a multiplicity of infection (MOI) of 1 per 100 μl of culture medium containing 10 μM of compounds (SCREEN-WELL Autophagy Library, Enzo Life Sciences) or 0.1% dimethyl sulfoxide (DMSO; control). The cells were fixed at 24 hours post-infection (hpi) by incubating with 150 μl of 3.7% formaldehyde for 30 min, rinsed with three times with PBS, then permeabilized with 50 μl of 0.5% Triton X-100 in PBS for 10 min. After three rinses with PBS, the cells were treated with 50 μl of 50 mM NH_4_Cl, rinsed three times with PBS, and incubated with 100 μl of Blocking One (Nacalai Tesque) for 1 h. Blocking One was replaced with 100 μl containing 1 μg/ml of anti-alphavirus mouse monoclonal IgG (Santa Cruz Biotechnology #3582; primary antibody) in 20-fold diluted Blocking One with PBS, then the cells were incubated at 4℃ overnight. The cells were rinsed three times with PBS, incubated at room temperature for 2 h in 100 μl of 2,000-fold diluted horseradish peroxidase (HRP)-conjugated-mouse IgG (Cell Signaling #7076; secondary antibody) in 20-fold diluted Blocking One with PBS, then rinsed three times with PBS. HRP signals were quantified using the luminescent substrate EzWestLumiOne (ATTO) and a Varioskan LUX microplate reader (Thermo Fisher Scientific).

To measure cell viability, cells were rinsed once with 100 μl of sterile water and stained with 30 μl of the Janus Green cell normalization stain (Abcam) at room temperature for 5 min. The cells were washed five times with sterile water, then incubated with 500 mM HCl (100 μl/well) for 10 min. The stain was eluted by shaking for 10 sec and transferred to clear, round-bottomed 96-well plates, then the optical density (OD) at 595 nm (OD_595_) was measured using the Varioskan LUX microplate reader.

### Antiviral activity assays

For the validation of candidate compounds, Vero cells (2 × 10^4^ cells/well) were seeded in 96-well plates, and the next day, the cells were incubated with 10 μM rotenone (Merck #8875), staurosporine (Abcam #ab120056), and anisomycin (Merck #A9789) for 1 h. Thereafter, the cells were infected with CHIKV SL11131 at an MOI of 1. At 2 hpi and cultured with 100 μl medium containing 10 μM compounds or 0.1% DMSO. The culture supernatants were collected at 24 hpi, and CHIKV titer in the culture supernatants were measured using plaque assays. The viability of infected Vero cells was assessed using CellTiter-Glo 2.0 Luminescent Cell Viability Assay (Promega) according to the manufacturer’s protocol.

To determine the effective inhibitory concentration of anisomycin, Vero (1 × 10^4^ cells/well in a 96-well plate one day prior to infection) and Huh7 (1 × 10^5^ cells/well in a 24-well plate one day prior to infection) cells were pretreated with 5-fold serial concentrations (64 pM – 1 μM) of anisomycin or 0.1% DMSO for 1 h. The cells were then infected with CHIKV at an MOI of 1. The culture medium was replaced at 2 hpi with a fresh medium containing the corresponding anisomycin concentrations. Culture supernatants were harvested two days post-infection (dpi) for plaque assays.

To determine the cytotoxic concentration, Vero and Huh7 cells were seeded in a 96-well plate (2.5 × 10^4^ cells/well) one day before treatment and cultured with 64 pM – 100 μM anisomycin. Cell viability was measured two days after treatment using alamarBlue Cell Viability Reagent (Thermo Fisher Scientific) according to the manufacturer’s protocol.

### Time-of-addition assay

For the pre-treatment experiment, Huh7 cells (1 × 10^5^ cells/well in 24-well plates at one day prior to assay) were incubated with 100 nM anisomycin for 1 h. The cells were washed twice with culture medium, then infected with CHIKV SL11131 at an MOI of 0.1. The culture medium was replaced at 2 hpi with fresh medium containing 0.1% DMSO. For the post-treatment experiment, Huh7 cells were infected with CHIKV at an MOI of 0.1 for 1 h, and after removal of the virus, the cells were cultured with fresh medium (0 h). Anisomycin (100 nM) was added to the cells at 0, 1, 2, 4, and 8 hpi. The controls were cells infected with CHIKV and treated with 0.1% DMSO. The viral titer in culture supernatants at 24 hpi was measured using plaque assays.

### Entry assay

Huh7 cells, which had been seeded in a 24-well plate at a density of 2 × 10^5^ cells/well one day before the assay, were infected with CHIKV at an MOI of 5 in the presence of 1 μM anisomycin or 0.1% DMSO at 37℃ for 1 h. Uninternalized virus particles were removed by washing the cells twice with cold PBS, followed by treatment with 0.25% trypsin-EDTA at 37℃ for 5 min. After three washes with cold PBS, total RNA was extracted from the cells using FastGene RNA Basic Kit (Nippon Genetics). Cell-associated viral RNA was detected by RT-qPCR using CHIKV-specific primers and a fluorescent probe [13] with iTaq Universal Probes One-Step Kit (Bio-Rad) and a QuantStudio 3 Real-Time PCR system (Thermo Fisher Scientific).

### Reporter replicon assay

To generate a CHIKV subgenomic RNA replicon expressing NanoLuc luciferase (Nluc), a gene unit containing puromycin resistance (pac), the foot-and-mouth disease virus 2A cleavage site (FMDV2A), and *Gaussia* luciferase (Gluc) genes in a CHIKV replicon plasmid (pT7/SL-Gluc) [13] was replaced with Nluc cDNA by inverse PCR using PrimeSTAR MAX DNA polymerase (Takara) and In-Fusion cloning (Clontech). A plasmid expressing a replication-defective reporter replicon, in which the catalytic motif in the RNA-dependent RNA polymerase (RdRp) of nsP4 was mutated, was similarly constructed by replacing the pac-FMDV2A-Gluc gene unit with Nluc cDNA in pT7/SL-Gluc-nsP4^GAA^ [14]. After confirming nucleotide sequence, the plasmid DNA was linearized with *Not*I, and the 5’-capped replicon RNA was transcribed *in vitro* using the mMESSAGE mMACHINE T7 Transcription Kit (Thermo Fisher Scientific), followed by purification using the FastGene RNA Basic Kit (Nippon Genetics Co., Ltd.).

Huh7 cells (2 × 10^4^ cells/well) were transfected with 200 ng of replicon RNA using 0.3 μl of Lipofectamine MessengerMAX (Thermo Fisher Scientific) in 96-well white plates (SPL Life Sciences) and cultured with 1 μM anisomycin or 0.1% DMSO on the following day. Two days after transfection, Nluc activity was measured using a Nano-Glo Live Cell Assay System (Promega) and the Varioskan Lux microplate reader.

### Entry-bypass assay

The plasmid pGEM-5Zf(+) containing a cDNA of CHIKV Ross genomic RNA (pT7-Ross [12]) was digested with *Not*I, and the 5’-capped full-length viral RNA was synthesized using the *Not*I-digested plasmid DNA and mMESSAGE mMACHINE T7 Transcription Kit. After purification using the FastGene RNA Basic Kit, *in vitro* transcribed RNA (5 μg) was transfected into 293T cells (2 × 10^5^ cells/well in 12-well plates one day before transfection) using Lipofectamine MessengerMAX. Eight hours after transfection, 1 μM anisomycin or 0.1% DMSO was added to the culture, and then the culture supernatant was collected 24 h after transfection to measure the infectious titer using plaque assays.

### Isolation and phenotyping of anisomycin-resistant virus

Huh7 cells (5 × 10^5^ cells/well in a 6-cm dish one day prior to infection) were infected with CHIKV SL11131 at an MOI of 1 and cultured in the presence of 100 nM anisomycin. Culture supernatant (3 ml) was harvested one week after infection, and a portion (100 μl) of the supernatant was added to a fresh Huh7 cell culture along with 200 nM anisomycin. This one-week infection and virus passage was repeated using 400, 800, and 1,600 nM of anisomycin. To confirm resistance to anisomycin, Huh7 cells were infected with the collected viruses at an MOI of 1 in the presence of 100 nM anisomycin, and viral titers in the culture supernatant at 2 dpi were measured using plaque assays.

To analyze the genome sequences of the resistant viruses, viral RNA was extracted from the harvested supernatant using the QIAamp Viral RNA Mini Kit (Qiagen) and reverse-transcribed using PrimeScript II 1st strand cDNA Synthesis Kit and random 6-mer primers (Takara Bio). Three DNA fragments covering the entire CHIKV genome were amplified by PCR using PrimeSTAR Max DNA Polymerase (Takara Bio) and the PCR primers listed in Supplementary Table 1. The PCR products were purified using FastGene Gel/PCR Extraction Kit (Nippon Genetics) and sequenced using the Sanger method (Eurofin Genomics) and the sequencing primers listed in Supplementary Table 1.

Point mutants found in resistant viruses by sequencing were introduced into pCMV-SL11131 by PCR-based site-directed mutagenesis using PrimeSTAR Max DNA Polymerase (Takara). The plasmids encoding the mutant CHIKV genome were confirmed by sequencing and used to transfect into 293T cells to produce infectious viruses. All plasmids were prepared using *Escherichia coli* strain DH5α (GMbiolab) and FastGENE Plasmid Mini (Nippon Genetics).

### Mouse model study

Animal studies proceeded in an Animal Biosafety Level 3 (ABSL3) facility according to the Guidelines for Animal Research of Osaka Medical and Pharmaceutical University, Faculty of Medicine. The Committee for Animal Use and Care of the Osaka Medical Pharmaceutical University approved experimental procedures (Approval no. AM24-030). Female C57BL/6J mice (4-week-old, purchased from Japan SLC) were subcutaneously infected with 100 PFU of CHIKV (in 20 μl PBS) on the dorsal side of the left hind foot under anesthesia with 2% isoflurane inhalation. At 3 hpi, the mice were intraperitoneally administered 150 μl of anisomycin (10 mg/kg in PBS containing 5% DMSO, n = 3) or control buffer (PBS containing 5% DMSO, n = 3) under anesthesia, followed by daily anisomycin administration at 4 dpi. Footpad joint swelling was measured daily from 0 to 7 dpi using a digital caliper. Blood (〜 100 μl) was sampled from the facial vein using a 5-mm animal lancet (Goldenrod) from 1 – 3 dpi, and serum was collected by centrifugation at 800 × *g* for 10 min. CHIKV titer in serum were determined using plaque assays.

### Statistical analysis

Data were statistically analyzed using one-way analysis of variance (ANOVA) with Dunnett’s multiple comparison tests, unless otherwise stated in the figure legends, using GraphPad Prism software. *P* values below 0.05 were considered statistically significant (*, *P* < 0.05; **, *P* < 0.01; ***, *P* < 0.001).

## Results

### Identification of anisomycin as a potential inhibitor of CHIKV replication

We used In-Cell ELISA (ICE) technique to screen small chemical compounds that could be potential CHIKV inhibitors (S1A Fig). In the In-Cell ELISA for CHIKV infection (hereafter referred to as ICE-CHIK), viral antigen expression in Vero cells infected with CHIKV SL11131 strain [12] was detected using anti-alphavirus primary and HRP-conjugated secondary antibodies, followed by measurement of the luminescent HRP signals (S1B Fig). After the detection of luminescent signals indicating viral replication, cell viability was measured by Janus Green staining, which indicates mitochondrial function (S1C Fig). We determined the robustness of ICE-CHIK to quantifying CHIKV infection using the Z’-factor, which was 0.59 (S2A Fig). The Z’-factor is a statistical parameter that indicates the quality of a high-throughput screen [15]. In addition, the OD_595_ of Janus Green staining was correlated closely with the number of cells seeded in 96-well plates (S2B Fig). Next, ICE-CHIK was used to evaluate the antiviral activity of mycophenolic acid (MPA), which has been reported to inhibit CHIKV *in vitro* [16]. We confirmed an MPA concentration-dependent reduction in the HRP signal (S2C Fig), with an EC_50_ of 232.6 ± 16.6 nM. HRP signals and Janus green staining were not affected by MPA (S2C and S2D Figs). These data indicated that ICE-CHIK established herein has a suitable hit window and reliability for screening CHIKV inhibitors.

Using ICE-CHIK, we performed a screening assay using a chemical library consisting of 94 compounds with autophagy-inducing or autophagy-inhibitory activities (SCREEN-WELL Autophagy Library, Enzo Life Sciences). The results of a primary screen using 10 μM compounds showed that average of HRP signals were reduced to 11.9%, 19.6%, 5.6%, 7.8%, 22.0%, 25.7%, 10.1%, 3.1%, and 9.2% in timosaponin AIII, rotenone, niclosamide, rottlerin, 17-AAG, geldanamycin, staurosporine, anisomycin, and cycloheximide-treated Vero cells, respectively, compared to that in 0.1% DMSO-treated control cells (Fig. 1A). Subsequent Janus Green staining showed that CHIKV-infected cell cultures treated with these compounds were almost 90% or more viable (Fig. 1A). A parallel ICE-CHIK screen of a chemical library containing 30 phosphatase inhibitors did not reveal any compound with anti-CHIKV activity that was not severely cytotoxic (S3 Fig).

**Fig. 1.**
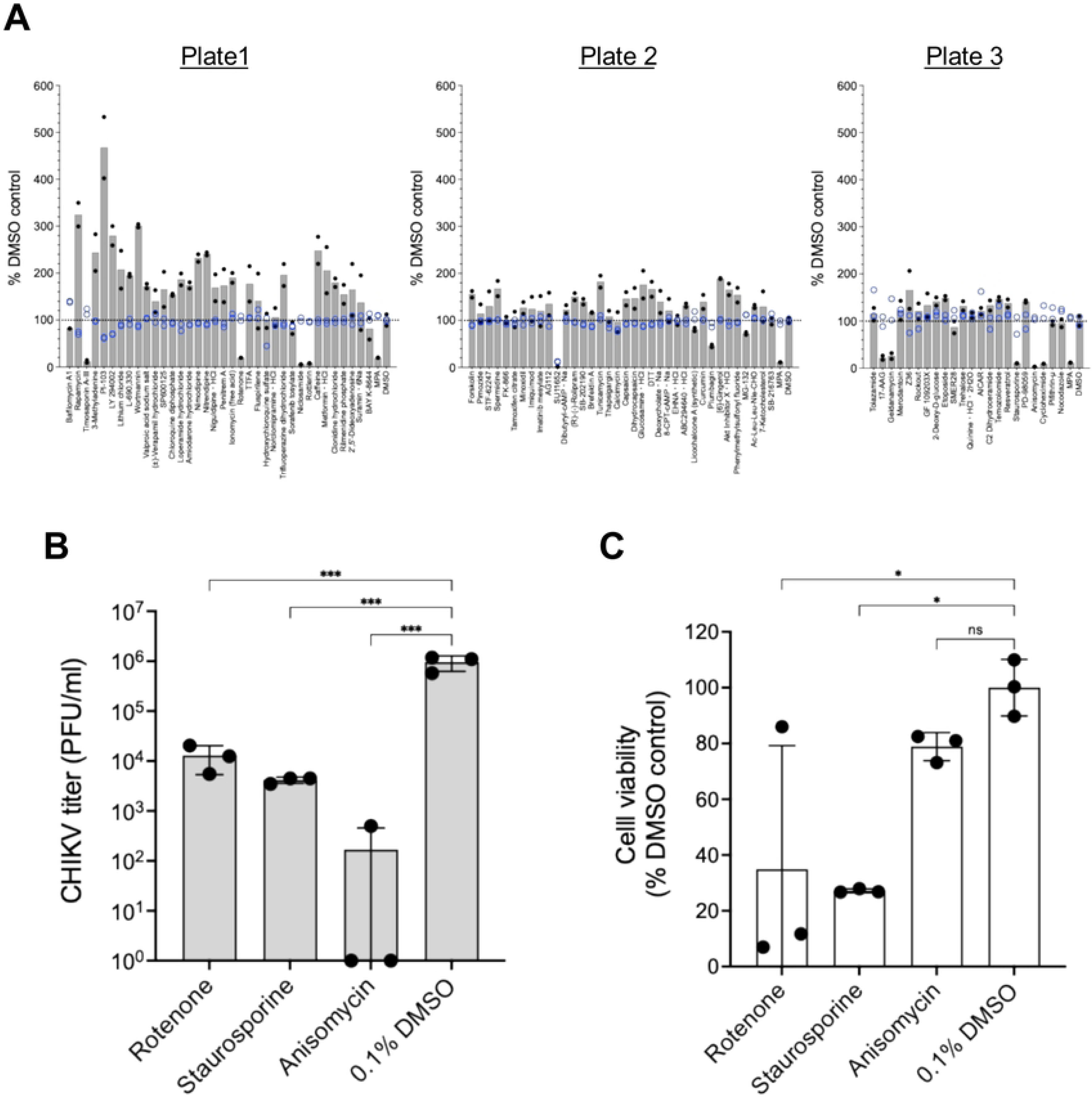
Identification and validation of anti-CHIKV compounds. (A) Primary screening of a chemical compound library using ICE-CHIK. Vero cells in 96-well plates (plates number 1, 2, and 3) were infected with CHIKV SL11131 at an MOI of 1 in the presence of 10 μM compounds or 0.1% DMSO. At 24 hpi, the cells were fixed, permeabilized, and incubated with an anti-alphavirus antibody, followed by an HRP-conjugated secondary antibody. HRP signals (black filled circles) were detected using a chemiluminescent substrate and quantified using a microplate reader. Cell viability was assessed by Janus Green staining and measuring OD_595_ (blue open circles). Results are expressed as relative ratios (%) compared to the DMSO-treated control. Gray bars and blue lines represent average HRP activity and Janus Green staining, respectively. (B) Validation of the candidate compounds. Vero cells were infected with CHIKV SL11131 at an MOI of 0.1, then incubated with 10 μM rotenone, staurosporine, and anisomycin or 0.1% DMSO. The viral titer in the culture supernatants at 2 dpi was measured using plaque assays (black circles). Average values of infectious titer (PFU/ml, gray bars) are shown with standard deviations (SD). (C) Cytotoxicity of candidate compounds. Vero cells were cultured with 10 μM selected compounds for 2 days, and cell viability was measured (black circles). Cell viability is expressed as relative mean ratios (%, white bars) with SD over that of DMSO-treated control cells.

Among the hit compounds, which included previously reported CHIKV inhibitors (niclosamide, rottlerin, 17-AAG, geldanamycin, and cycloheximide [17–21]), rottlerin, staurosporine, and anisomycin were further validated. Vero cells were infected with CHIKV SL11131 at an MOI of 1 and incubated with 10 μM of the candidate compounds. At 2 dpi, CHIKV titers in the culture supernatant were measured using plaque assays. Anisomycin had the strongest inhibitory effect on viral replication (Fig. 1B) without a significant loss of cell viability (Fig. 1C). Therefore, anisomycin was selected to further analyze its anti-CHIKV activity.

### Evaluation of anti-CHIKV activity of anisomycin

Anisomycin (Fig. 2A) is a pyrrolidine-type antibiotic that was originally isolated from *Streptomyces griseolus* [22], and it inhibits protein synthesis in eukaryotic and prokaryotic cells [23]. Anisomycin is also shown to inhibit encephalomyocarditis virus (EMCV [24]), Japanese encephalitis virus (JEV [25]), dengue virus (DENV [26]), Zika virus (ZIKV [26]), and Coxsackievirus B (CVB [27]). However, the anti-CHIKV activity of anisomycin has remained unknown. Anisomycin dose-dependently reduced the viral titer in culture supernatants of Vero cells infected with CHIKV SL11131 (Fig. 2B), which is classified as an IOL [28]. This led to a 50% effective concentration (EC_50_) of 0.215 ± 0.092 nM, a 50% cytotoxic concentration (CC_50_) in mock-infected Vero cells (Fig. 2C) of 390.6 ± 141.1 nM, and the selective index (SI) was > 1,800. Anisomycin also dose-dependently inhibited CHIKV SL11131 in the human hepatoma Huh7 cells (Fig. 2D), with an EC_50_ of 4.8 ± 2.1 nM. The EC_50_ against Ross strain, which was first isolated in 1953 and classified as an ECSA lineage [12], in Huh7 cells was 4.3 ± 2.4 nM (Fig. 2E). In Huh7 cells, the CC_50_ of anisomycin was 85.4 ± 26.0 nM (Fig. 2F) and the range of providing SI values was 17.7 – 19.8.

**Fig. 2.**
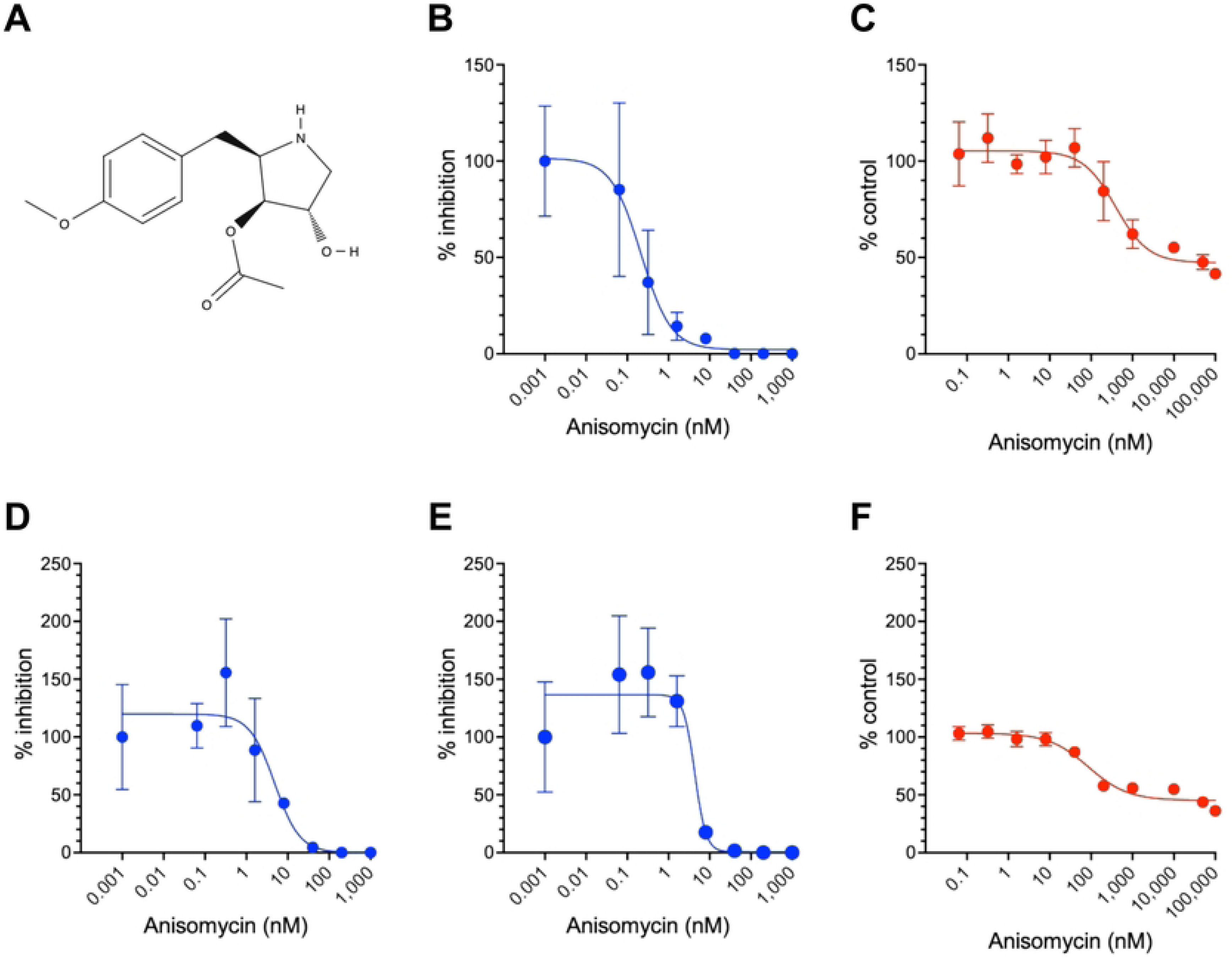
Anti-CHIKV activity of anisomycin. (A) Chemical structure of anisomycin. (B, D, E) Dose-dependent inhibition of CHIKV by anisomycin. Vero (B) and Huh7 (D, E) cells were cultured for 1 h with 64 pM – 1 μM anisomycin or 0.1% DMSO (control), then infected with CHIKV SL11131 (B, D) or Ross (E) strain at an MOI of 0.1. After infection, cells were incubated with various concentrations of anisomycin (or 0.1% DMSO), and the viral titer in culture supernatants harvested at 2 dpi was determined using plaque assays. The average % inhibition relative to the viral titer in the DMSO control is presented with SD. (C, F) Determination of cytotoxic concentration. Vero (C) and Huh7 (F) cells were cultured with 64 pM – 1 μM anisomycin or 0.1% DMSO for two days, and cell viability was analyzed. Values are expressed as relative mean % with SD over that of DMSO controls.

### Replication step of CHIKV inhibited by anisomycin

To estimate the stage of CHIKV replication at which anisomycin inhibited viral replication, a time-of-addition assay was conducted. Anisomycin (100 nM) was added to Huh7 cells before or after CHIKV infection, and the viral titer in the culture supernatants harvested at 24 hpi was measured using plaque assays. Adding anisomycin before CHIKV infection did not inhibit CHIKV replication in Huh7 cells (Fig. 3, pre-treatment). However, starting anisomycin treatment immediately after the 1 h exposure of the virus significantly reduced the CHIKV titer of the culture supernatant at 24 hpi (Fig. 3, post-treatment, 0 h). Although significant inhibition of CHIKV replication was observed even at 8 h post-treatment, the antiviral activity of anisomycin diminished when added at 4 hpi and beyond (Fig. 3).

**Fig. 3.**
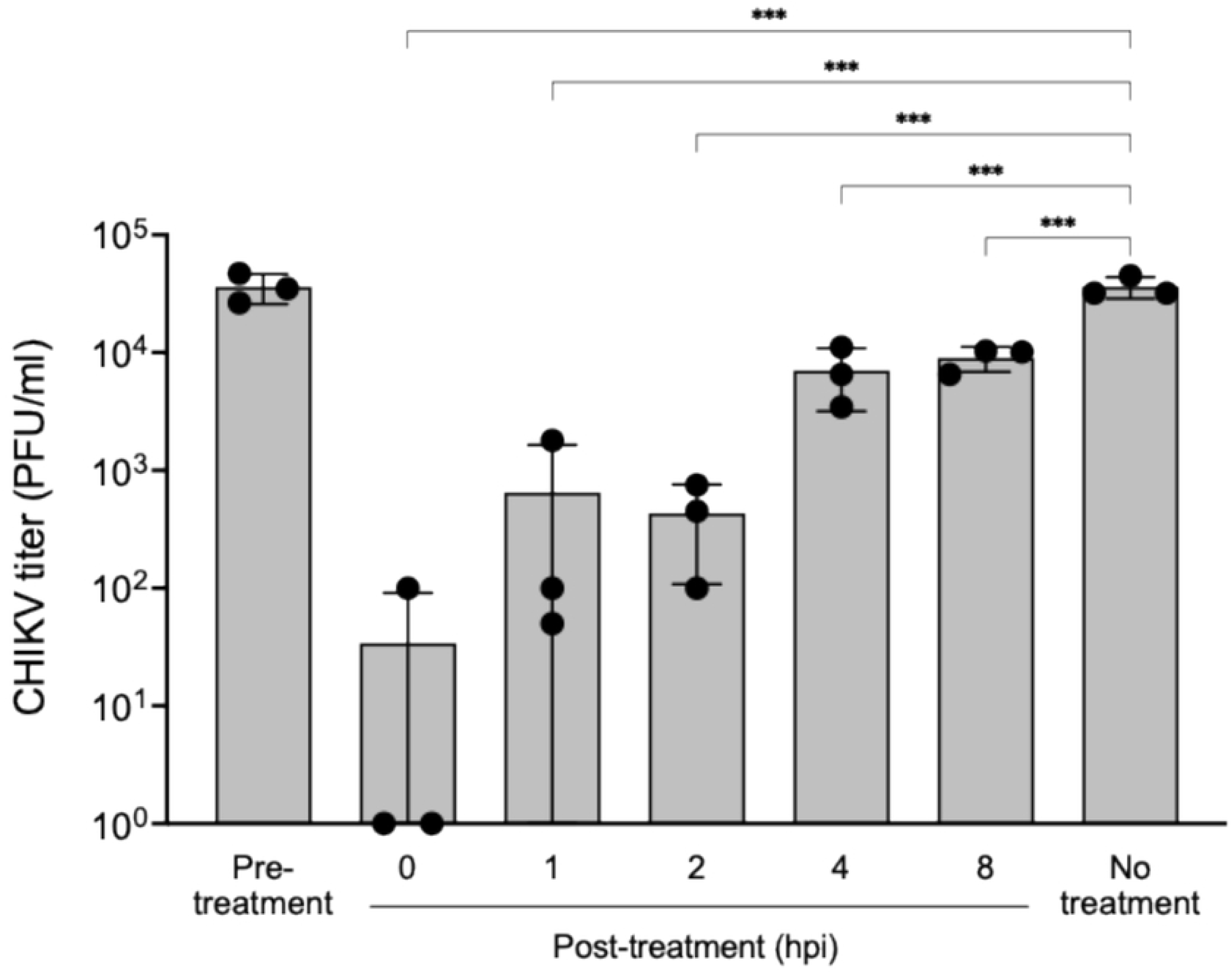
Time-of-addition assay. Huh7 cells were initially incubated with 100 nM anisomycin for 1 h, washed, then infected with CHIKV SL11131 at an MOI of 0.1 (pre-treatment). For post-treatment experiments, cells were infected with CHIKV for 1 h, then 100 nM anisomycin (or 0.1% DMSO) was added at 0, 1, 2, 4, and 8 hpi. Culture supernatants were harvested at 24 hpi, and viral titers (PFU/ml) were measured using plaque assays (black circles). Gray bars represent average viral titers with SD.

To clarify the effect of anisomycin on CHIKV attachment and entry steps, an RT-qPCR-based virus entry assay was performed [14]. Huh7 cells were exposed to CHIKV in the presence of anisomycin at 37℃ for 1 h, which allowed virion binding and internalization. Uninternalized viruses were removed with trypsin, and the cells were then washed with cold PBS. The RT-qPCR results with CHIKV-specific primers revealed that the same levels of cell-associated CHIKV RNA were detected in anisomycin- and control (DMSO)-treated cells (Fig. 4A).

**Fig. 4.**
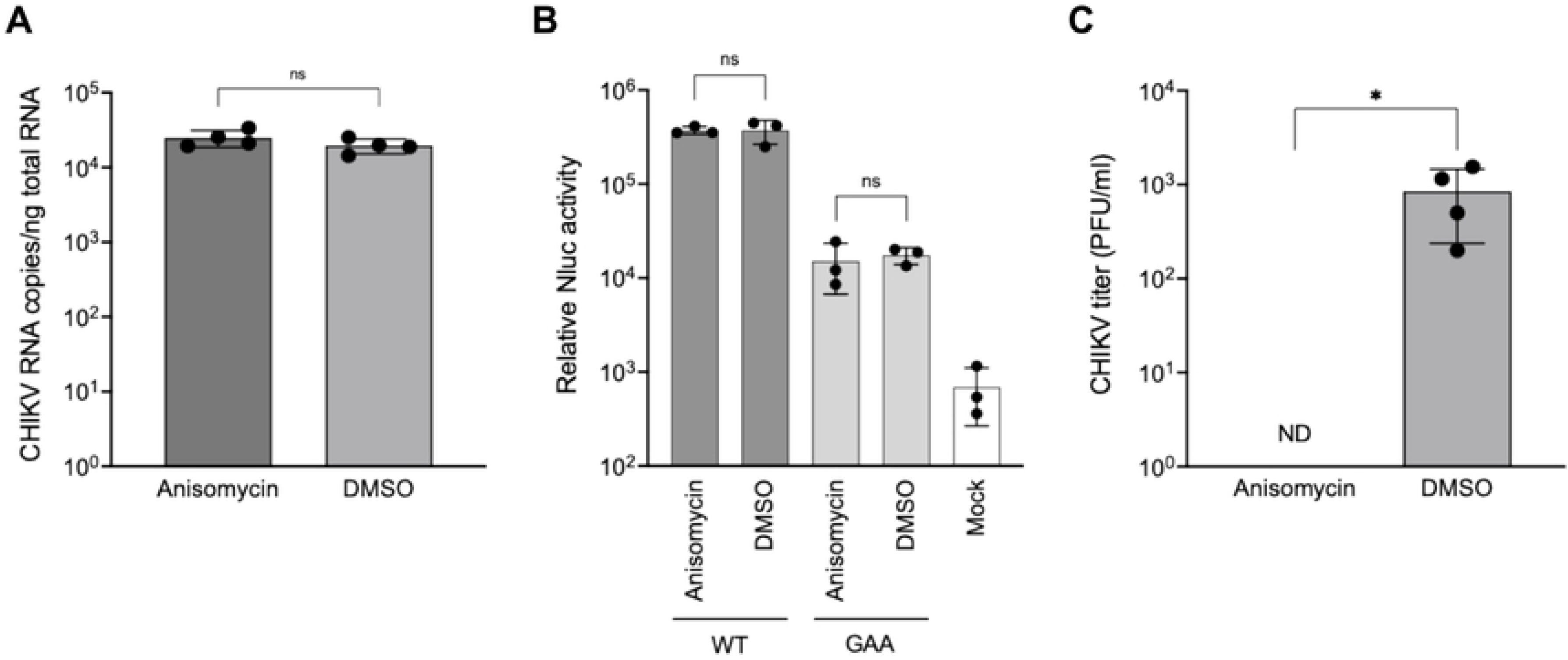
Inhibition of infectious CHIKV production stage by anisomycin. (A) Entry assay. Huh7 cells were exposed to CHIKV SL11131 at an MOI of 5 for 1 h at 37℃ in the presence of 1 μM anisomycin or 0.1% DMSO. After removing uninternalized virus by PBS washes and trypsin-EDTA treatment, total RNA was extracted and viral RNA was quantified by RT-qPCR using CHIKV-specific primers and a fluorescent probe (black circles). (B) Reporter replicon assay. The 5’-capped and 3’-polyadenylated RNA of the CHIKV subgenomic replicon expressing Nluc (WT) or its replication-defective mutant that harbors a mutated catalytic motif in nsP4 (GAA) was synthesized by *in vitro* transcription and transfected into Huh7 cells. Cells were cultured with 1 μM anisomycin or 0.1% DMSO, then Nluc activity was measured two days after transfection (black circles). Mock represents background luminescent responses of non-transfected cells. (C) Entry-bypass assay. Full-length CHIKV RNA was synthesized by *in vitro* transcription and transfected into 293T cells. After 8 h, the cells were incubated with 1 μM anisomycin or 0.1% DMSO, then culture supernatants were harvested 24 h after transfection. CHIKV titers (PFU/ml) were measured using plaque assays (black filled circles). ND, below the detection limit of the plaque assay (5 PFU/ml). (A, B, C) All bars represent mean values with SD. Statistical significances were determined by the unpaired *t*-tests with Welch’s correction (A and B) or the Mann-Whitney U test (C). ns, not significant.

Next, to examine whether anisomycin affected CHIKV RNA replication and translation steps, *in vitro* transcribed RNA of a CHIKV subgenomic replicon expressing Nluc was transfected into Huh7 cells, followed by incubation with 1 μM anisomycin. Measurement of Nluc activity 2 days after transfection showed that anisomycin did not affect reporter replicon activity compared to the control condition (Fig. 4B, wild type [WT]). Although anisomycin is reported to inhibit protein translation [23], the Nluc activity of a replication-defective CHIKV replicon, in which the GDD motif in the active site of nsP4 RdRp was changed to GAA [29], did not differ between the cells incubated with anisomycin and DMSO control (Fig. 4B, GAA).

The effect of anisomycin on post-entry events of CHIKV was further evaluated using entry-bypass assay, in which viral genomic RNA is directly transfected into cells to skip virion binding and entry processes [30]. 293T cells were transfected with CHIKV RNA, which had been transcribed from plasmid DNA containing the full-length viral cDNA *in vitro* [12], then treated with 1 μM anisomycin at 8 h after transfection. A plaque assay to measure infectious virus titers in the culture supernatants at 24 h after transfection revealed a significant reduction in the CHIKV titer after anisomycin treatment (Fig. 4C). Taken together, these data indicated that anisomycin acts on the processes that lead to the production of infectious CHIKV after viral RNA synthesis and protein translation.

### Characterization of anisomycin-resistant virus

To investigate the CHIKV gene that is directly or indirectly affected by anisomycin, a resistant variant selection experiment was performed by passaging on CHIKV-infected Huh7 cells with increasing concentrations of anisomycin (100 to 1,600 nM, Fig. 5A). When the final supernatant collected after four rounds of passages (P5 virus, Fig. 5A) was used, efficient CHIKV replication was observed in a new Huh7 cell culture even in the presence of 100 nM anisomycin (Fig. 5B). Sanger sequencing of cDNA from the P5 virus RNA using the sequencing primers (Supplemental Table 1) revealed four changes compared to the original CHIKV SL11131 cDNA sequence in pCMV-SL11131 [12]. The anisomycin-resistant virus harbored a G to A change at nucleotide position 540 (G540A), T1446C, G4425A, and T4644G, which led to a Gly to Thr change at amino acid position 155 in nsP1 (nsP1 A155T), Ser to Pro change at position 457 in nsP1 (S457P), Gly to Arg at position 117 in nsP3 (G117R), and Tyr to Asp at position 190 in nsP3 (Y190D), respectively (Fig. 5C).

**Fig. 5.**
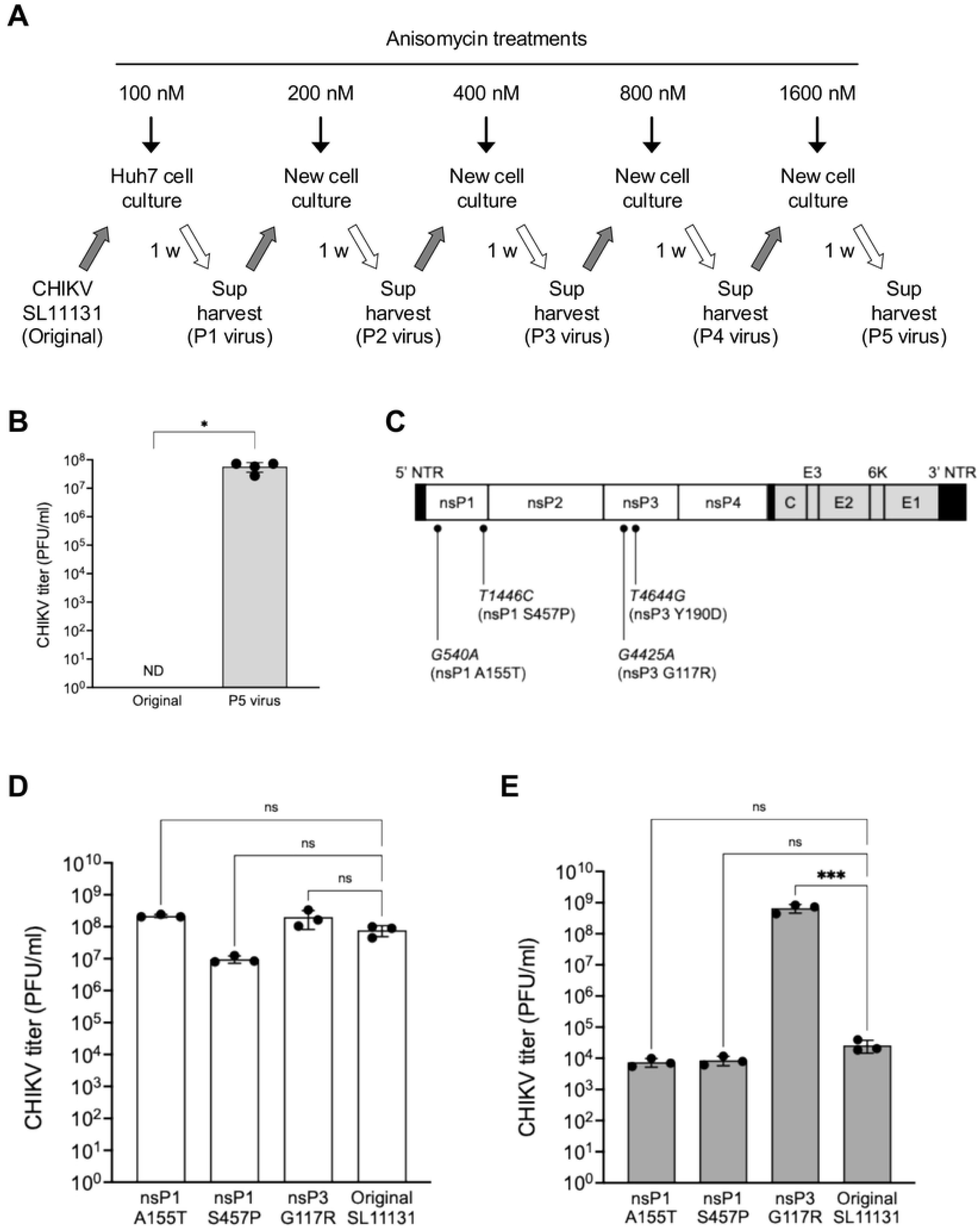
Characterization of anisomycin-resistant CHIKV. (A) Scheme for selecting anisomycin-resistant mutants. Huh7 cells were infected with CHIKV SL11131 and cultured with anisomycin (100 nM). Culture supernatant (sup) was harvested one week after infection and added to a new Huh7 cell culture with higher concentrations of anisomycin (200 nM). This one-week infection and virus passage were repeated with increasing concentration (400, 800, and 1,600 nM) of anisomycin. (B) Analysis of resistance by selected viruses against anisomycin. Huh7 cells were infected with the final resistant CHIKV (P5 virus) or with the original CHIKV SL11131 and cultured with 100 nM anisomycin. Viral titers (PFU/ml, black circles) were determined using a plaque assay. Bars represent all mean values with SD. ND, below the detection limit of the plaque assay (< 5 PFU/ml). Statistical significance was determined by the Mann-Whitney U test. (C) Mutations found in the anisomycin-resistant CHIKV. Viral RNA was purified from the P5 virus stock and subjected to RT-PCR to synthesize cDNA. Fragments of DNA covering the entire CHIKV genome were amplified by PCR and analyzed using Sanger sequencing. (D, E) Replication efficiency of point mutant CHIKV. Mutations found in anisomycin-resistant CHIKV were individually introduced into pCMV-SL11131, and infectious viruses of each point mutant, except for nsP3 Y190D, were produced by plasmid transfection in 293T cells. Mutant (nsP1 A115T, nsP1 S457P, and nsP3 G117R) and original CHIKV were used to infect Vero cells without (D) or with anisomycin (E). Viral titers (PFU/ml, black circles) were determined by plaque assays. Bars represent all mean values with SD. ns, not significant.

To determine the mutation(s) that contribute to phenotypic resistance to anisomycin, the identified mutations were individually introduced into pCMV-SL11131. Although the introduction of the Y190D mutation into SL11131 cDNA did not produce infectious viruses according to the results of plaque assays, sufficient infectious titers of virus were obtained by plasmid transfection of other single mutants, as well as the original pCMV-SL11131. In the absence of anisomycin treatment, the replication abilities of nsP1 A155T, nsP1 S457P, and nsP3 G117R mutant viruses were comparable to those of the original virus in Vero cells (Fig. 5D). However, the nsP1 A155T and S457P viruses were sensitive to the antiviral effect of 100 nM anisomycin, as in the original SL11131, whereas the nsP3 G117R mutant was insensitive (Fig. 5E), indicating that the Gly-to-Arg change at position 117 in nsP3 was primarily responsible for resistance to anisomycin.

### Evaluation of anti-CHIKV activity of anisomycin *in vivo*

We assessed the efficacy of anisomycin in a mouse model *in vivo* [31]. C57BL/6J mice (n = 3) were subcutaneously inoculated with CHIKV in the left hindfoot, followed by daily intraperitoneal anisomycin administration from the day of infection to 4 dpi. Swelling of the inoculated foot was monitored daily for 7 days, and the CHIKV titer in sera from 1 – 3 dpi was measured using plaque assays (Fig. 6A). In our mouse model experiment, a derivative of CHIKV SL11131 designated SL-Large, in which an Arg at the amino acid position 55 in E2 was replaced with a Gly (E2 R55G, S4A Fig [32]), was used because a preliminary study revealed that SL-Large had better replication and more foot swelling activity in C57BL/6 mice than the original CHIKV SL11131 (manuscript in preparation). The susceptibility of SL-Large to anisomycin was confirmed *in vitro* in Huh7 cells, with an EC_50_ of 11.2 ± 2.0 nM (S4B Fig).

**Fig. 6.**
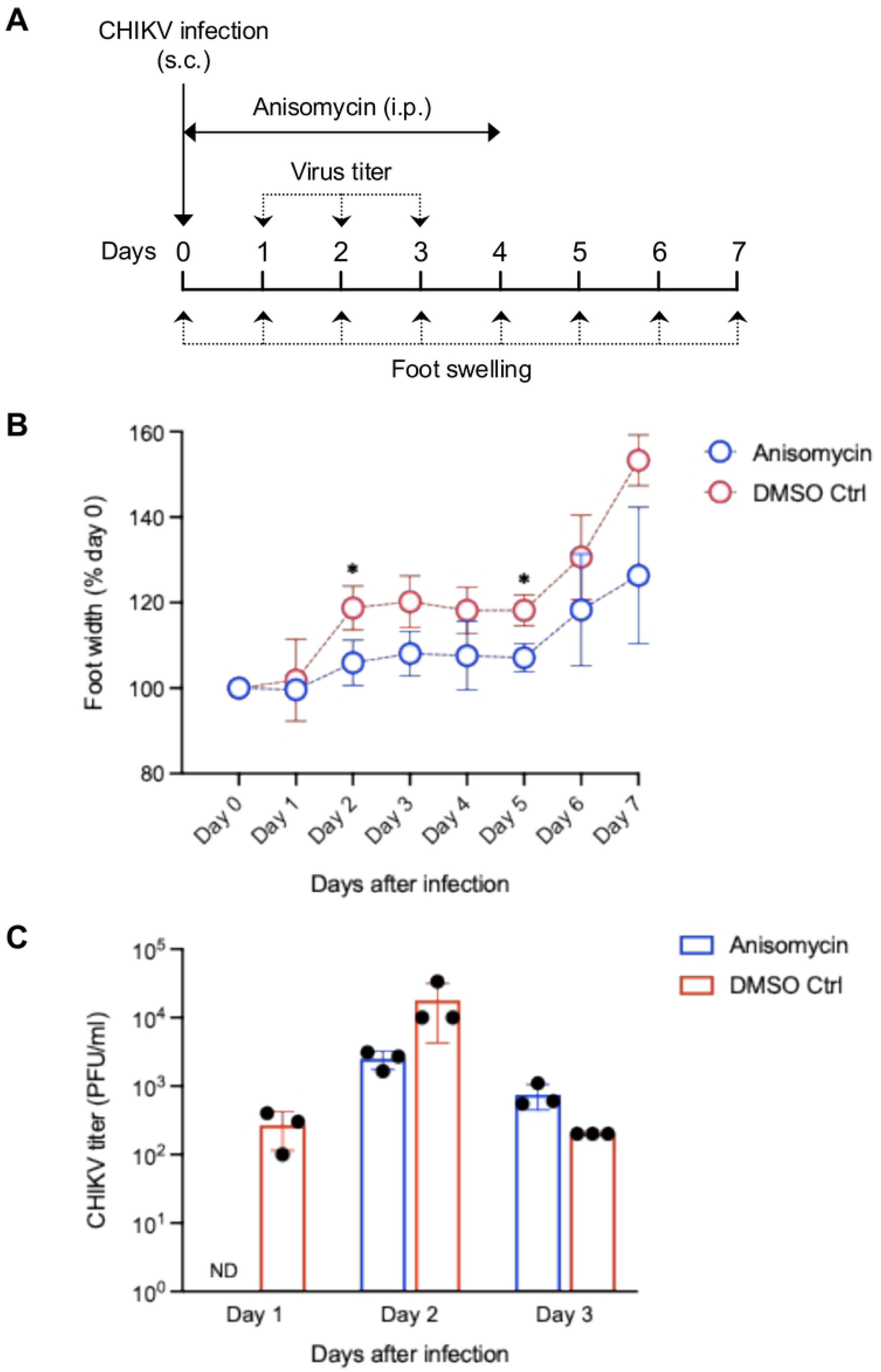
Effects of anisomycin against CHIKV infection *in vivo*. (A) Experimental design for assessing anisomycin effects in mouse models infected with CHIKV. The left footpads of female C57BL/6J mice (*n* = 3 per treatment) were subcutaneously infected (s.c.) with 100 PFU of CHIKV. Anisomycin (10 mg/kg) or control buffer was then injected intraperitoneally (i.p.) at 3 hpi, followed by daily administration until 4 dpi. Swollen footpad joints were measured daily from 0 to 7 dpi. Blood was collected from the facial vein from 1–3 dpi, and CHIKV titers in the serum samples were quantified using plaque assays. (B) Anisomycin ameliorated CHIKV-induced footpad swelling. Time course of footpad swelling measured daily for 7 dpi in mice treated with anisomycin (blue circles) or control buffer (red circles). Data are presented as relative mean ratios (%) at 0 dpi with SD (*n* = 3 per treatment). Statistical significance was determined by the Mann-Whitney U test. (C) Effects of anisomycin on viremia. CHIKV titers in blood sera collected from 1 – 3 dpi were determined using plaque assays (PFU/ml, black circles). Blue (anisomycin-treated) and red (control buffer-treated) bars indicate average viral titer in model mice. ND, below the detection limit of the plaque assay (< 50 PFU/ml).

Footpad swelling induced by CHIKV SL-Large infection continued to increase in mice treated with anisomycin and DMSO for 7 dpi *in vivo*, whereas it was improved in mice treated with anisomycin-treated mice (Fig. 6B). Measurement of CHIKV titer in the sera showed that the peak viremia level was lower in anisomycin-treated mice, although an increased virus titer in the anisomycin-treated mice was observed at 3 dpi (Fig. 6C). These findings showed that anisomycin alleviated foot swelling induced by CHIKV and reduced viremia *in vivo*.

## Discussion

Once regarded as a disease endemic to East Africa, CHIKF has emerged as a global viral infection that has spread to numerous countries in Africa, Asia, Europe, and the Americas. The US FDA recently approved the live attenuated vaccine VLA1553 (IXCHIQ) as a preventive measure against CHIKV infection [11]. Despite this, CHIKF treatment still depends on symptom management, highlighting the ongoing challenge in developing effective therapeutics. The present study identified anisomycin as a potent CHIKV inhibitor that probably interferes with the production of infectious viruses, with nsP3 being a proposed target viral protein. Anisomycin notably decreased CHIKV pathogenicity in mouse models.

Anisomycin was originally isolated from *Streptomyces* bacteria as an antibiotic that was toxic to specific protozoa and yeasts [23]. However, it also acts against picornaviruses and flaviviruses [24–27], suggesting broad-spectrum inhibitory activity. Anisomycin blocks protein translation in ribosomes [23]. In line with this activity, it inhibits protein synthesis in both EMCV and ZIKV [24,26]. In contrast, anisomycin did not affect reporter protein expression in our investigation using a replication-defective CHIKV replicon RNA expressing the Nluc (Fig. 4B). This indicated that anisomycin did not inhibit CHIKV during viral translation. Low concentrations of anisomycin have been reported to stimulate multiple mitogen-activated protein kinase (MAPK) pathways, including p38 and c-Jun N-terminal kinases (JNK) [33,34]. However, anisomycin-induced CHIKV inhibition in Huh7 cells was not affected by the p38 inhibitor SB202190 (S5A Fig) or the MEK1/2 inhibitor U0126 (S5B Fig). Furthermore, the anti-CHIKV activity of anisomycin was enhanced by a specific JNK inhibitor SP600125 that is shown to suppresses CHIKV infection [35] (S5C Fig). Anisomycin is also reported to inhibit CVB replication *via* the proteasomal degradation of eukaryotic translation elongation factor 1 alpha 1 (eEF1A1), which is required for CVB replication [27]. However, because the siRNA-mediated depletion of eEF1A1 [36] did not affect CHIKV replication (S6 Fig), anisomycin was unlikely to suppress CHIKV replication *via* eEF1A1 degradation. Interestingly, the entry bypass assay revealed that a significant reduction in the infectious titer of CHIKV produced from transfected viral RNA was observed by anisomycin treatment (Fig. 4C). These results suggest that anisomycin acts as an inhibitor during infectious virion production step after CHIKV protein translation.

In this study, we developed anisomycin-resistant CHIKV to enhance understanding of its mechanism of inhibition and identified four mutations in the nsP1 and nsP3 regions of resistant viruses (Fig. 5C). Introducing individual mutations into the original CHIKV infectious clone indicated that the substituted G117R mutation in nsP3 conferred significant resistance to anisomycin (Fig. 5E). Kinetic analysis of the sequencing data of P1–P5 viruses revealed that the G117R mutation was already extant in the P3 virus population and became dominant in the P4 and P5 populations (S7. Fig), supporting the importance of this amino acid residue to anisomycin susceptibility. Although nsP3 is the least understood protein among those encoded by the CHIKV genome, it is a component of the RNA replicase complex and comprises an N-terminal macrodomain, a central alpha unique domain, and a C-terminal hypervariable region [37]. Among the three domains, the anisomycin-resistance mutation G117R is located in the macrodomain. Macrodomain is widely found in eukaryotes and microorganisms and is also encoded by several positive-strand RNA viruses. It possesses enzymatic activity that catalyzes hydrolysis to remove adenosine diphosphate (ADP) ribose from proteins [38]. Macrodomain hydrolytic activity plays a critical role in CHIKV replication and pathogenesis [39]. A previous study of a large plaque phenotype obtained by the continuous passage of a primary CHIKV isolate found that the G117R mutation was required not only to form large plaques but also for pathogenicity in suckling mice, suggesting that this mutation in nsP3 promotes the replication efficiency of CHIKV [40]. Our findings showed that anisomycin probably inhibited the post-translational steps of CHIKV infection (Fig. 4). Although nsP3 is thought to be primarily involved at the viral RNA replication stage, it would also be involved in CHIKV particle formation through an unidentified mechanism, and anisomycin might hamper the function of nsP3 at the late stage of viral replication. A recent structural study revealed that CHIKV nsP3 assembles into helical tubular structures in the cytoplasm containing viral RNA and capsid proteins, suggesting its involvement in nucleocapsid formation and infectious virion assembly [41]. Additionally, some α-mangostin-derived synthetic compounds that bind to the nsP3 macrodomain was shown to inhibit the late stages of CHIKV replication [42]. Whether anisomycin binds directly to nsP3, or affects nsP3 *via* other factors that interact with nsP3, or influences the function of nsP3 remains to be determined. However, our results suggest that this could serve as a foundation for the development of new inhibitors that can suppress viral particle formation *via* nsP3.

In addition to the antiviral activity *in vitro*, we evaluated the efficacy of anisomycin in CHIKV-infected C57BL/6J mice *in vivo*. The results showed that daily administration of anisomycin reduced joint swelling in feet inoculated with the CHIKV (Fig. 6B). Although anisomycin did not decrease viremia levels throughout the measurement period, it suppressed CHIKV infection in mice at 1 and 2 dpi (Fig. 6C). Thus, the findings *in vivo* suggest that anisomycin contributes to controlling the CHIKV burden and mitigating disease severity to some extent. However, the strong anti-CHIKV activity of anisomycin *in vitro* was not evident in our mouse models *in vivo*. One plausible reason would be that a sufficient amount of anisomycin might not have reached the joints and muscle tissues, where CHIKV preferentially replicates. Therefore, more effective methods are needed to administer anisomycin, and pharmacokinetics in CHIKV-infected mouse models require future elucidation.

## Conclusions

Our study revealed that anisomycin is a potent CHIKV inhibitor and that it relieved virus-induced foot swelling in mice. Anisomycin likely blocks infectious virus production, and the susceptibility of CHIKV to anisomycin is determined by the nsP3 macrodomain. Given its inhibitory activity against other RNA viruses, anisomycin is a promising lead compound for the development of broad-spectrum antiviral drugs.

## Acknowledgments

We thank Dr. Atsushi Tanala (Osaka Medical and Pharmaceutical University) for his technical advice in mouse model study. This work was supported by the Research Program on Emerging and Re-emerging Infectious Diseases of the Japan Agency for Medical Research and Development (Grant No. JP25fk0108656).

## Supporting information

**S1 Fig. Overview of ICE-CHIK.** (A) Vero cells seeded in a white 96-well plate were infected with CHIKV SL11131 at an MOI of 1 and incubated with 10 μM compounds or 0.1% DMSO (day 0). At 24 hpi, the cells were fixed, permeabilized, and blocked before incubation with anti-alphavirus primary antibody overnight at 4℃ (day 1). The cells were washed, then HRP-conjugated secondary antibody was applied, and signals were detected using a chemiluminescent substrate and a microplate reader (day 2, CHIKV replication). Cell viability was assessed in the same plate by staining with Janus Green and measuring OD_595_ (day 2, cytotoxicity). (B) The luminescent signal from HRP in 96-well plates, containing CHIKV-infected cell cultures treated with different compounds in individual wells, were visualized using a FUSION FX7 imaging system (Vilber). Images of representative areas in 96-well plates are shown. (C) Wells identical to those in (B) were stained with Janus Green after HRP detection, and the stained images were captured using a flatbed scanner.

**S2 Fig. Evaluation of ICE-CHIK.** (A) In-Cell ELISA of CHIKV-infected (positive control; blue) and mock-infected (negative control; red) Vero cells was performed in 25 wells of a 96-well plate. Activities of HRP are expressed as relative light units (RLU, open circles). Solid lines represent means of positive and negative controls, and dashed lines represent three standard deviations of each control that were used to determine the Z’-factor. (B) Vero cells were seeded onto a 96-well plate at cell counts decreased by half of 20,000 cells, and the Janus Green staining of ICE-CHIK was performed. Linear regression was analyzed using absorbance (X-axis) and OD_595_ data (Y-axis). (C, D) ICE-CHIK was performed using CHIKV-infected Vero cells (Blue) in the presence of 2-fold serial dilutions of MPA starting with 40 μM or 0.1% DMSO. To examine the effects of MPA treatment on HRP activity and Janus Green staining, the same In-Cell ELISA was performed without CHIKV infection (red). Inhibition curves were generated from three independent experiments by a four-parameter fitting method using GraphPad Prism software. (C) HRP activity and (D) Janus Green staining data.

**S3 Fig. ICE-CHIKV screening using a phosphatase inhibitor library.** (A) ICE-CHIK was performed in the presence of 10 μM compounds (SCREEN-WELL Phosphatase Inhibitor Library, Enzo Life Sciences) or 0.1% DMSO. Black filled circles, HRP activity; gray bars, average of three measurements of HRP activity; blue open circles, Janus Green staining. Data are expressed as percentages relative to the DMSO-treated control.

**S4 Fig. Features of CHIKV SL-Large.** (A) SL-Large contains an Arg (R) to Gly (G) substitution at amino acid position 55 of E2. (B) Susceptibility of SL-Large to anisomycin. Vero cells were infected with SL-Large at an MOI of 0.1 and incubated with 5-fold serial dilutions of anisomycin starting with 10 μM or 0.1% DMSO. Virus titers in culture supernatants harvested at 2 dpi were determined using plaque assays. Average inhibition ratios (%) versus the viral titer in the DMSO control treatment is presented with SD (*n* = 3).

**S5 Fig. Effects of MAPK inhibitors on the anti-CHIKV activity of anisomycin.** (A, B, C) Huh7 cells (1 × 10^5^ cells/well) seeded in 24-well plates one day before infection were incubated with anisomycin (10 nM) alone, MAPK inhibitor alone (SB202190 [10 nM], U0126 [20 μM], or SP600125 [10 μM]), or a combination of anisomycin with one of the MAPK inhibitors for 1 h and infected with CHIKV SL11131 at an MOI of 1, followed by cultivation under the corresponding treatment conditions. The virus titer in the culture supernatants at 2 dpi was measured using plaque assays (black circles). (A, B, C) All bars represent mean values with SD. ns, not significant.

**S6 Fig. Effects of eEF1A1 depletion on CHIKV replication.** Huh7 cells (1 × 10^5^ cells) were reverse-transfected with 5 pmol of siRNA duplex against eEF1A1 (sieEF1A1), CHIKV nsP3 (sinsP3), or nonspecific siRNA (siCtrl) using Lipofectamine RNAiMAX Transfection Reagent (Thermo Fisher Scientific). The next day, the cells were infected with CHIKV SL11131 at an MOI of 0.1, and the viral titer in the culture supernatants at 2 dpi was measured using plaque assays (black circles). Gray bars represent mean values with SD. ns, not significant.

**S7 Fig. Sequencing chromatograms of CHIKV during anisomycin-resistant mutant selection.** Changes in nucleotides and amino acid residues of P1 – P5 viruses are shown.

